# Tracheophyte genomes keep track of the deep evolution of the *Caulimoviridae*

**DOI:** 10.1101/158972

**Authors:** Seydina Diop, Andrew D.W. Geering, Françoise Alfama-Depauw, Mikaël Loaec, Pierre-Yves Teycheney, Florian Maumus

## Abstract

Endogenous viral elements (EVEs) are viral sequences that are integrated in the nuclear genomes of their hosts and are signatures of viral infections that may have occurred millions of years ago. The study of EVEs, coined paleovirology, provides important insights into virus evolution. The *Caulimoviridae* is the most common group of EVEs in plants, although their presence has often been overlooked in plant genome studies. We have refined methods for the identification of caulimovirid EVEs and interrogated the genomes of a broad diversity of plant taxa, from algae to advanced flowering plants. Evidence is provided that almost every vascular plant (tracheophyte), including the most primitive taxa (clubmosses, ferns and gymnosperms) contains caulimovirid EVEs, many of which represent previously unrecognized evolutionary branches. In angiosperms, EVEs from at least one and as many as five different caulimovirid genera were frequently detected, and florendoviruses were the most widely distributed, followed by petuviruses. From the analysis of the distribution of different caulimovirid genera within different plant species, we propose a working evolutionary scenario in which this family of viruses emerged at latest during Devonian era (approx. 320 million years ago) followed by vertical transmission and by several cross-division host swaps.

## Introduction

Although the field of viral metagenomics is rapidly expanding the repertoire of viral genome sequences available for evolutionary studies ^1^, it only provides a picture of viral diversity over a very short geological time scale. However, viruses can leave molecular records in the genomes of their hosts in the form of endogenous viral elements (EVEs). EVEs are viral sequences that have been inserted in the nuclear genomes of their hosts by either active or passive integration mechanisms and in many cases have been retained for extended periods of time, sometimes millions of years. The study of EVEs does allow the evolution of viruses to be traced, much like a fossil record ^2^. For example, the study of endogenous retroviruses has uncovered the extended diversity and host range of retroviruses, and has provided evidence that they have a marine origin, and that they developed in parallel with their vertebrate hosts more than 450 million years ago (MYA; ^3 4 5^).

Plant EVEs were first discovered a little more than 20 years ago ^6^ but have only received a fraction of the research attention directed towards endogenous retroviruses in humans and other animals. Most characterized plant EVEs are derivatives of viruses in the family *Caulimoviridae* ^7^. The *Caulimoviridae* is one of the five families of reverse-transcribing viruses or virus-like retrotransposons that occur in eukaryotes ^8^, and is the only family of viruses with a double-stranded DNA genome that infects plants (https://talk.ictvonline.org/). Unlike retroviruses, members of the *Caulimoviridae* do not integrate in the genome of their host to complete their replication cycle. However, caulimovirid DNA can occasionally integrate passively into their host genome. In fact, five of the eight officially recognized genera of the *Caulimoviridae* have EVE counterparts in at least one plant genome ^7 9^.

Recently, Geering et al. ^10^ showed that EVEs from an additional tentative genus of the *Caulimoviridae*, called ‘Florendovirus’, are widespread in the genomes of cultivated and wild angiosperms, and provided evidence for the oldest EVE integration event yet reported in plants, at 1.8 MYA ^10^. Furthermore, sister taxa relationships between florendoviruses in South American and Australian plants suggested Gondwanan links and a minimum age of 34 MYA for this virus group based on estimates of when land bridges between these two continents were severed. About 65% of all angiosperm species that were examined contained endogenous florendoviruses and for five, these sequences contributed more than 0.5% of the total plant genome content. Furthermore, the discovery of endogenous florendoviruses in basal ANITA (*Amborella*, *Nymphaeales* and *Illiciales*, *Trimeniaceae*-*Austrobaileya*) grade angiosperm species also showed that beyond mesangiosperms, the host range of the *Caulimoviridae* extends or once extended to the most primitive known angiosperms. Furthermore, some reconstructed endogenous florendovirus genomes were bipartite in organization, a genome arrangement that is unique among viral retroelements ^10^. Overall, the work of Geering et al. (2014) demonstrated that analyzing the genetic footprints left by viruses in plant genomes can contribute to a better understanding of the long-term evolution of the *Caulimoviridae*.

In this study, we hypothesized that other unrecognized groups of caulimovirid EVEs could exist, particularly in some of the more primitive plant taxa that have recently been subject to plant genome sequencing initiatives. Following an extensive search in the genomes of over 70 plant species, we discovered EVEs from several novel genera. We show that the *Caulimoviridae* host range extends throughout the Euphyllophyte and Lycopodiophyte clades, which constitute the Tracheophyta, and surpasses that of any other plant virus family. By analyzing the distribution of different genera of the *Caulimoviridae* within different plant species, we unveil a complex pattern of associations and propose a scenario in which the *Caulimoviridae* would have emerged approximately 320 million years ago.

## Results

### Augmenting the diversity of known endogenous caulimovirids

The reverse transcriptase (RT) domain is the most conserved domain in the genome of viral retroelements and is used for classification ^11 12^. The strong sequence conservation of this domain allows high quality alignments to be generated, even for distantly related taxa. We have thus used a collection of RT domains from known exogenous and endogenous caulimovirids to search for related sequences across the breadth of the Viridiplantae (four green algae, one moss, one lycopod, four gymnosperms, and 62 angiosperms; Supplementary Table 1) using tBLASTn.

Initially, over 8,400 protein-coding sequences were retrieved, all containing an RT domain with a best reciprocal hit against members of the *Caulimoviridae*, as opposed to the closely related *Metaviridae* (Ty3/Gypsy group LTR retrotransposons). To provide a preliminary classification, sequences with at least 55% amino acid identity to each other were clustered and then iteratively added to our reference set of RT domains to build a sequence similarity network. The successive networks were examined manually and representative sequences from each cluster were kept only when creating substantially divergent branches so as to cover an extended diversity of caulimovirid RT with a core sequence assortment. While this network-based approach cannot be taken as phylogenetic reconstruction, it provided a practical method to explore diversity.

In the final sequence similarity network (Figure 1), 17 groups with deep connections were identified, hereafter referred to as operational taxonomic units (OTUs). Remarkably, nine of these OTUs were distinct from recognized genera of the *Caulimoviridae*. Four of these novel OTUs were exclusively composed of sequences from gymnosperms, thereby representing a new and significant host range extension for the *Caulimoviridae*. These OTUs were named Gymnendovirus 1 to 4. Two other novel OTUs were composed of RTs from various angiosperms and were named Xendovirus and Yendovirus. The last three novel OTUs were lesser populated, comprising sequences from one or two plant species (*Petunia inflata* and *Petunia axillaris*; *Vitis vinifera*; *Glycine max*; named species-wise: Petunia-, Vitis-, and Glycine-endovirus). This initial search therefore enabled uncovering a significantly augmented diversity of caulimovirid RTs.

**Figure 1.**
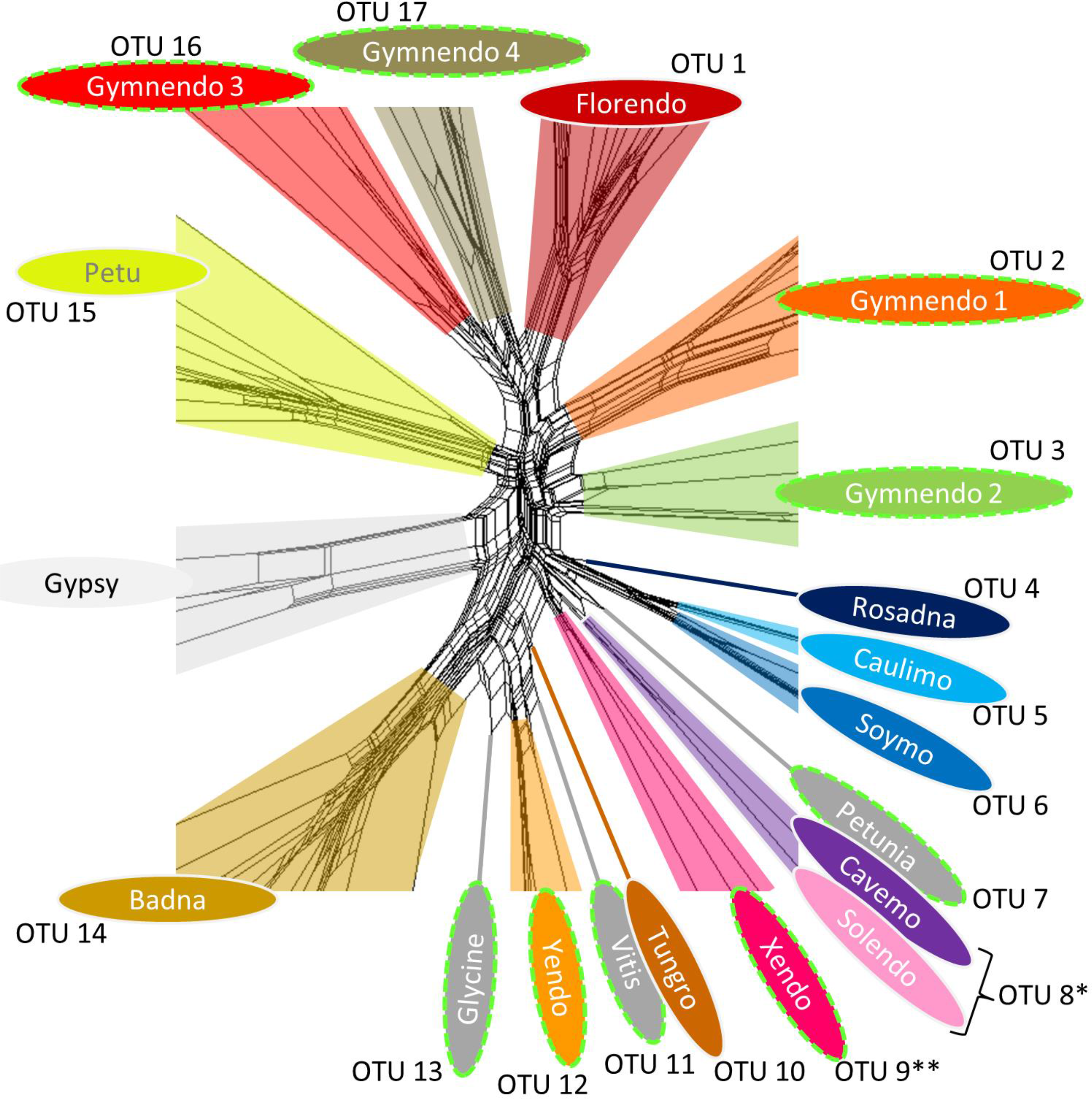
Augmented diversity of the *Caulimoviridae*. Core of a phylogenetic network constructed using an alignment of amino acid reverse transcriptase (RT) sequences from reference genera, representative endogenous caulimovirid RTs (ECRTs) and Ty3/Gypsy LTR retrotransposons. The full network is available in Supplementary Figure 1. This representation allows determining 17 Caulimoviridae OTUs. OTU names have dashed lime green outline when they include no known reference genera (referred to as novel OTUs). Each fill color corresponds to a different OTU except for OTUs comprising only a representative ECRT sequence that are colored with dark grey and named after the only host plant genome they were detected in at this stage (Petunia-, Vitis-, and Glycine- virus). * RT clustering at 55% identity groups *Cavemovirus* and *Solendovirus* into a single OTU (OTU 8). ** Sequences grouped in the Xendovirus OTU appeared to be paraphyletic after phylogenetic reconstruction (see Figure 3).

### Endogenous caulimovirid RT (ECRT) density across the Viridiplantae

To perform a more comprehensive search for ECRTs in our collection of plant genomes, we used the sequences from the final phylogenetic network (Figure 1) to search for ECRT nucleotide sequences that do not necessarily retain uninterrupted open reading frames. Using tBLASTn, we detected 14,895 genomic loci representing high-confidence ECRT candidates. Remarkably, ECRTs were found in nearly all seed plants, ranging from gymnosperms (ginkgo and conifers) to angiosperms. Quantitatively, over one-thousand ECRTs were detected in the genome assemblies of the gymnosperms *Picea glauca* (white spruce) and *Pinus taeda* (loblolly pine), as well as from the solanaceous plant species *Capsicum annuum* (bell pepper) (Figure 2A). In general, we observed a positive correlation between plant genome size and the number of ECRTs, although there were notable exceptions, such as the monocot *Zea mays* (maize), which has a relatively large genome at 2.1 Gb but no detectable ECRT. Five other seed plants from our sample also lacked ECRTs, including two other monocots (*Zostera marina* and *Oryza brachyantha*) and three dicots in the order *Brassicales* (*Arabidopsis thaliana*, *Schrenkiella parvula* and *Carica papaya*). When the number of ECRTs was normalized against genome size, *Citrus sinensis* (sweet orange) and *Ricinus communis* (castor bean) had the highest densities at 2.3 and 2 ECRTs per Mb, respectively (Figure 2B). The primitive ANITA grade angiosperm *Amborella trichopoda* also had a relatively high density of ECRTs (1 ECRT per Mb) compared to an average density of 0.2 ECRT per Mb across the 62 seed plant species that were examined.

**Figure 2.**
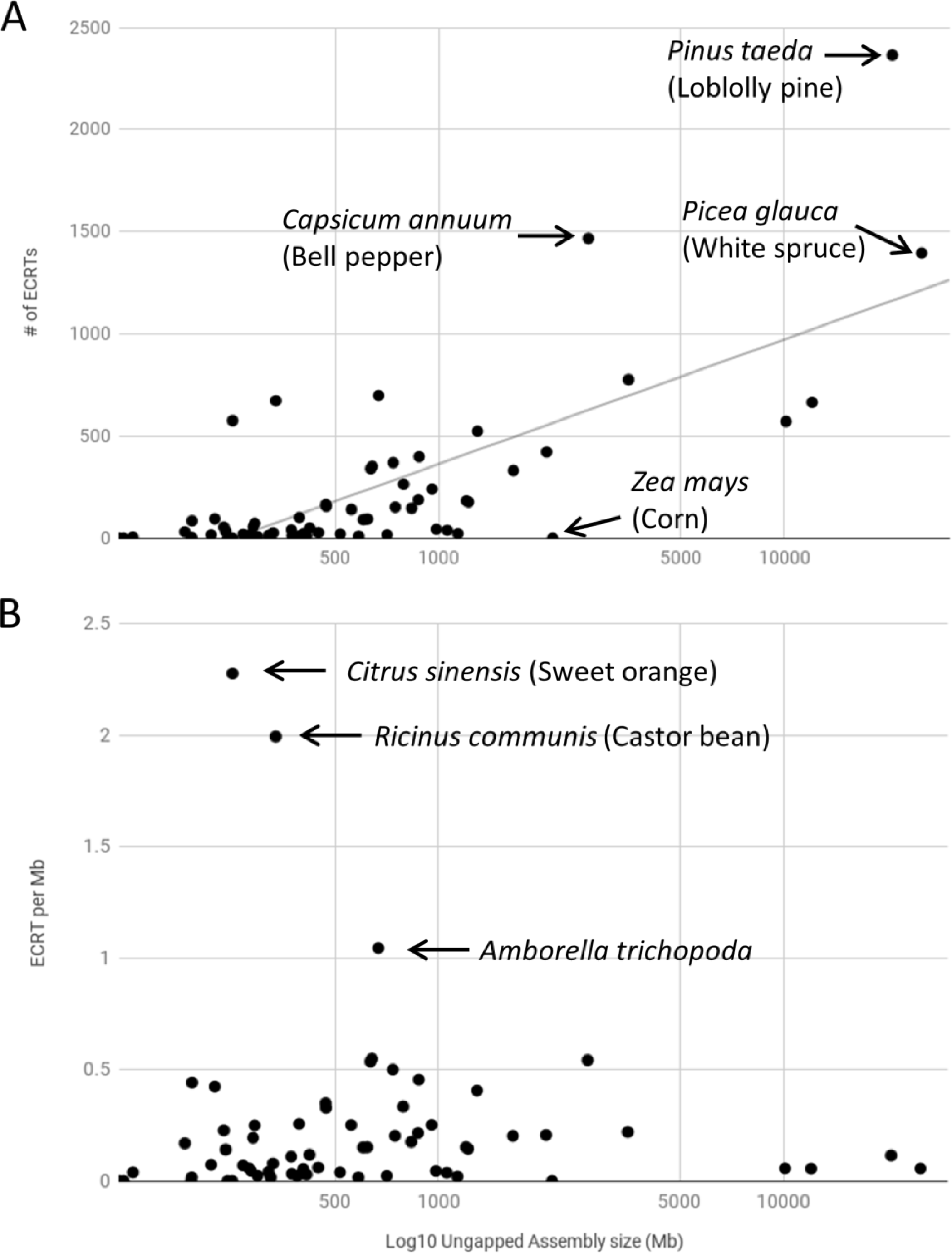
Highly variable ECRT numbers and density across plants. (A) Number of ECRTs found in each plant genome as function of Log10 genome size expressed in megabases (assembly gaps excluded). Logarhitmic trendline indicates moderate correlation between the number of ECRT and genome size (R^2^=0.544). (B) Density of ECRTs per megabase in each plant genome as function of Log10 genome size expressed in megabases (assembly gaps excluded). In (A) and (B), arrows indicate a sample of outlier dots and the corresponding plant species name.

### Caulimovirid sequences also detected in ferns and a clubmoss

From the plant genomes examined thus far, ECRTs were detected in gymnosperm genomes but not in those from the spikemoss *Selaginella moellendorffii* and from the moss *Physcomitrella patens*, which belong to the more basal land plant divisions Lycophyta and Bryophyta, respectively. Ferns (class Polypodiopsida) represent a bifurcation between Lycophyta and seed plants in the evolution of the Viridiplantae ^13^, but no high quality genome assemblies are publicly available for these plants. However, six fern genomes have recently been sequenced at low coverage (approximately 0.4 to 2 x genome size equivalent ^14^ and we therefore screened these datasets for the presence of ECRTs. A total of twenty-one protein-coding ECRTs were detected in genomic contigs from five of the six fern species examined (Supplementary Table 1). Sequence similarity network reconstruction using representative fern ECRTs revealed that they form two novel OTUs that were named Fernendovirus 1 and 2 and numbered OTU #18 and 19, respectively (Supplementary Figure 2).

Additional basal lineages of the Viridiplantae are represented in the 1,000 plant transcriptomes generated by the 1KP initiative ^15 16^. From this dataset, we found two transcript contigs (2.4 and 2.8 kilobases long, respectively) in the fern *Botrypus virginianus* (identifier BEGM-2004510) and *Lindsaea linearis* (identifier NOKI-2097008), which contained ECRTs (Supplementary file 1). Remarkably, we identified one more transcript contig (identified as ENQF-2084799, 2kb) that contained an ECRT in the clubmoss *Lycopodium annotinum*, which belongs to the *Lycopoda*, the most basal radiation of vascular plants (Tracheophyta). It is not possible to determine whether the mRNAs were transcribed from exogenous viruses or from EVEs.

### Phylogenetic reconstruction

Complete or near complete viral genomes were reconstructed from each novel OTU except Fernendovirus 1 and 2 (Supplementary file 1). From the fern genomic data sets, we were able to reconstruct fragments of Fernendovirus 1 and 2 genomes that contain sufficient genome coverage for phylogenetic analysis. We also used the complete genomes of the type species of the eight currently recognized genera in the family *Caulimoviridae* (*Badnavirus*, *Caulimovirus*, *Cavemovirus*, *Petuvirus*, *Rosadnavirus*, *Solendovirus*, *Soymovirus* and *Tungrovirus*), those of two unassigned viruses, Blueberry fruit drop-associated virus (BFDaV, ^17^) and Rudbeckia flower distortion virus (RuFDV, ^18^), and caulimovirid EVEs from the tentative genera Orendovirus ^19^ and Florendovirus (Geering et al., 2014). From this library of sequences, we aligned conserved protease, reverse transcriptase and ribonuclease H1 domains to build a maximum likelihood phylogenetic tree (Figure 3). Importantly, all newly identified EVEs grouped within the *Caulimoviridae* with strong bootstrap support.

**Figure 3.**
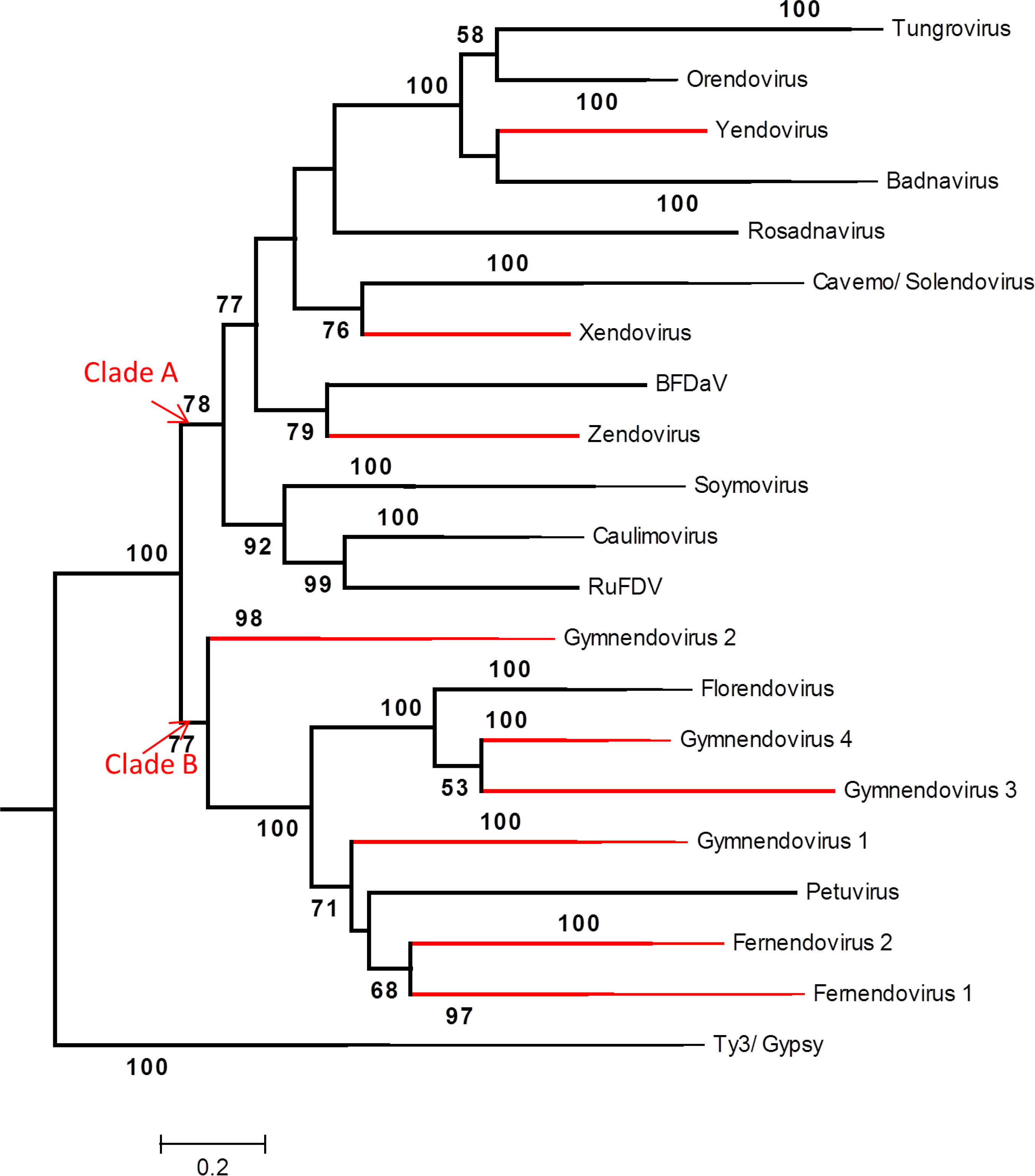
Phylogeny of the *Caulimoviridae*. Phylogenetic tree obtained by maximum likelihood search from a multiple sequence alignment of the genomic regions containing protease, reverse transcriptase and ribonuclease H1 domains from known (black) and novel (red) Caulimoviridae genera. The sequences from Ty3/Gypsy LTR retrotransposons are used as outgroups. Bootstrap support values below 50% are not shown. Sequences from members of the novel genera are available in supplementary data. Closely related sequences were collapsed into branches. The sequences contained in each branch are as follows. Orendovirus: Aegilops tauschii virus (AtV), Brachypodium distachyon virus (BdV); *Tungrovirus*: *Rice tungro bacilliform virus* (RTBV), Rice tungro bacilliform virus isolate west Bengal (RTBV); *Badnavirus*: *Commelina yellow mottle virus* (ComYMV), *Banana streak OL virus* (BSOLV); Yendovirus: Capiscum annuum virus; Zendovirus: Fragaria vesca virus; Blueberry: Blueberry fruit drop associated virus (BFDaV); *Caulimovirus*: *Cauliflower mosaic virus* (CaMV), *Figwort mosaic virus* (FMV); Rudbeckia: Rudbeckia flower distortion virus (RuFDV); *Soymovirus*: *Soybean chlorotic mottle virus* (SoyCaulimoviridae), *Peanut chlorotic streak virus* (PCSV); *Solendovirus*: *Sweet potato vein clearing virus* (SPVCV), *Tobacco vein clearing virus* (TVCV); *Cavemovirus*: *Cassava vein mosaic virus* (CsVMV), *Sweet potato collusive virus* (SPCV); *Petuvirus*: *Petunia vein clearing virus* (PVCV); *Rosadnavirus*: *Rose yellow vein virus* (RYVV); Florendovirus: Fragaria vesca virus (FvesV), Mimulus guttatus virus (MgutV); Gymnendovirus 1: Pinus taeda Gymnendovirus 1, Picea glauca Gymnendovirus 1; Gymnendovirus 2: Pinus taeda Gymnendovirus 2, Picea glauca Gymnendovirus 2, Ginkgo biloba Gymnendovirus 2; Gymnendovirus 3: Pinus taeda Gymnendovirus 3; Gymnendovirus 4: Pinus taeda Gymnendovirus 4, Picea glauca Gymnendovirus 4; Fernendovirus 1: Cystopteris protrusa Fernendovirus 1 contig 1, and the transcript scaffolds BEGM- 2004510 from *Botrypus virginianus*, NOKI-2097008 from *Lindsaea linearis*, and ENQF-2084799 from *Lycopodium annotinum*; Fernendovirus 2: Dipteris conjugata Fernendovirus 2 Contigs 2, 4 and 1319.

In agreement with previous studies ^10^, the tree revealed two sister clades, hereafter referred to as clade A and B. Clade A comprised sequences from representatives of Xendovirus and Yendovirus OTUs and from members of the genera *Caulimovirus*, *Soymovirus*, *Rosadnavirus*, *Solendovirus*, *Cavemovirus*, *Badnavirus*, *Tungrovirus* and Orendovirus, as well as RuFDV and BFDaV. The Xendovirus OTU was found to be polyphyletic, hence a new taxon, Zendovirus, was raised to include the EVE from *Fragaria vesca*, while the EVE from *Gossypium raimondii* (cotton) was retained in Xendovirus. The Yendovirus OTU, comprising the EVE from *Capsicum annuum* (bell pepper), fell in the subclade comprising bacilliform-shaped viruses in the genera *Badnavirus* and *Tungrovirus*. The reconstructed genomes from novel OTUs found in single dicot species (Petunia-, Vitis-, and Glycine-endovirus) were discarded from the phylogenetic reconstruction as they significantly weakened the robustness of the tree. However, they unambiguously fell within clade A (data not shown).

Clade B comprised EVEs from the four gymnendovirus OTUs, the two fernendovirus OTUs, as well as representatives of the genus *Petuvirus* and the tentative genus Florendovirus. Gymnendovirus 2 EVEs were sister to all other clade B viruses, indicating that this group of viruses arose in gymnosperms. Interestingly, the angiosperm-infecting caulimovirids in this clade were polyphyletic, indicating independent origins and probable large host jumps of the most recent common ancestors. Fernendovirus 1 and 2 were monophyletic, and the club moss EVE placed within Fernendovirus 1. Again, the fernendoviruses appear to have arisen after a large host range swap of the most recent common ancestor.

### ECRT distribution across seed plant genomes

To address the distribution of caulimovirid EVEs in our collection of plant genomes, we determined the most likely position within the reference *Caulimoviridae* phylogenetic tree proposed above (Figure 3) for the 14,895 ECRTs that we collected from seed plant genomes using the pplacer program ^20^. For this, we extracted ECRT loci extending upstream and downstream so as to retrieve potential sequences containing the contiguous fragment corresponding to the protease, RT and ribonuclease H1 domains. Using more relaxed length criteria, we extracted a total of 134 ECRT loci from the fern genomic data set that we also attempted to place on our reference tree.

Applying this strategy, we were able to assign unambiguous phylogenetic position on specific OTUs to a total of 13,834 ECRTs (Figure 4), the remaining ECRT loci being placed on inner nodes of the reference tree. Overall, we observed striking differences between *Caulimoviridae* genera for both the number of ECRT loci and the number of plant species in which they were found. For instance, Florendovirus ECRT loci were the most abundant, amounting to an overall total of 5,000 copies, and they were also found in the highest number of host species (46 of the 62 seed plant species that were screened). *Petuvirus* ECRT loci were also well represented, with an overall total of 1,900 copies found in a total of 27/62 seed plant species, especially in dicots. Among the novel OTUs, ECRTs classified as Yendovirus were found in the largest number of species, including monocots and dicots (Figure 4).

**Figure 4.**
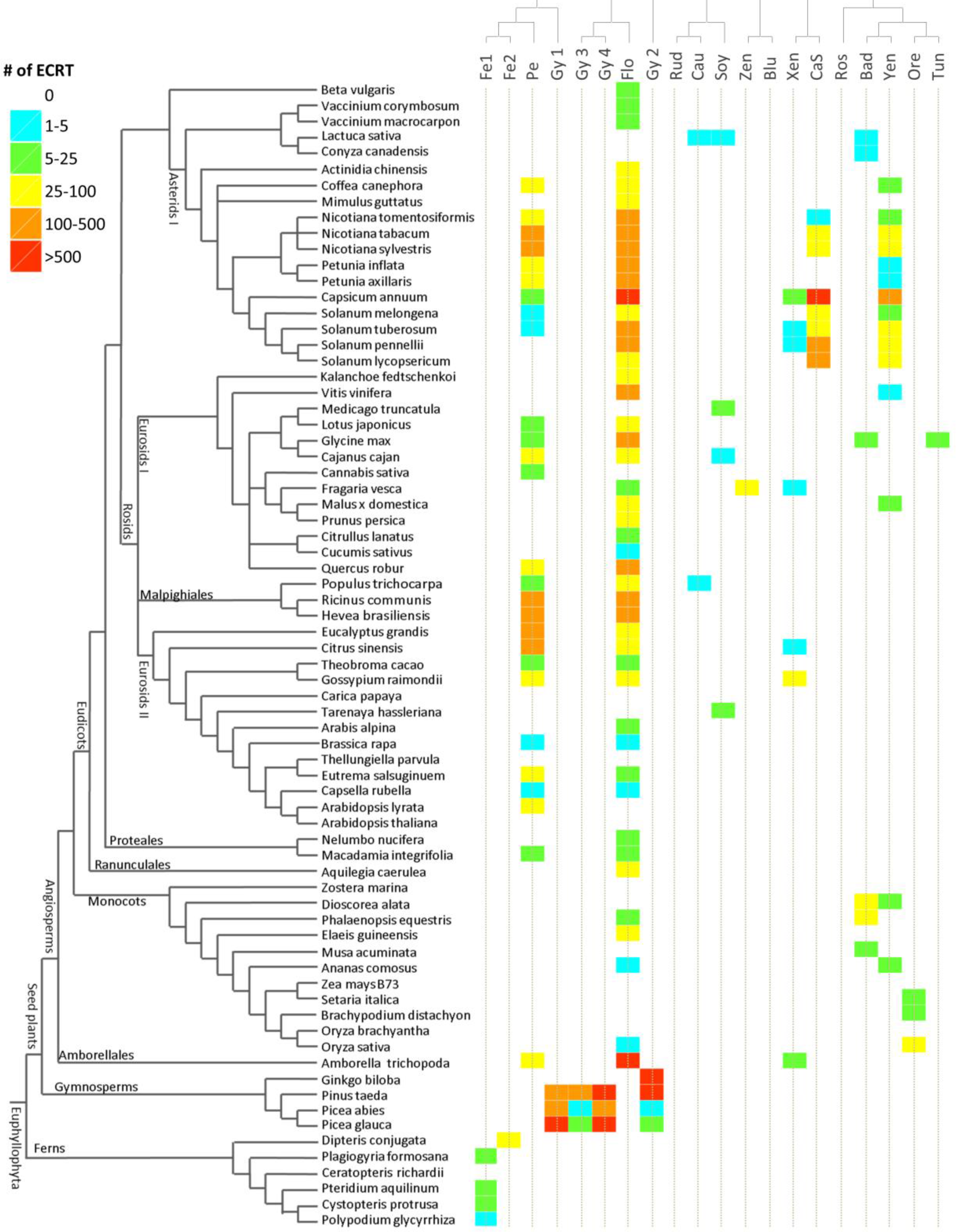
Distribution of caulimovirid EVEs in Euphyllophyte. The left tree represents a cladogram of Euphyllophyte species investigated in this study. The name of major branches and nodes is indicated. The top tree represents the topology of the phylogenetic tree obtained in Figure 3. At the intersection of these two trees, color code indicates the number of ECRT loci classified into each Caulimoviridae genus for each plant species. Abbreviations of virus genera are as follows: Pe (Petuvirus), Gy1 (Gymnendovirus 1), Gy2 (Gymnendovirus 2), Gy3 (Gymnendovirus 3), Gy4 (Gymnendovirus 4), Fe1 (Fernendovirus 1), Fe2 (Fernendovirus 2), Flo (Florendovirus), Soy (*Soymovirus*), Rud (Rudbeckia flower distortion virus), Cau (*Caulimovirus*), Blu (Blueberry fruit drop- associated virus), Zen (Zendovirus), Xen (Xendovirus), Yen (Yendovirus), CaS (*Cavemovirus* +*Solendovirus*), Ros (Rosadnavirus), Bad (*Badnavirus*), Tun (*Tungrovirus*), Ore (Orendovirus).

Most importantly, the detailed distribution of *Caulimoviridae* ECRTs in plant genomes reveals striking differences between ferns, gymnosperms and angiosperms (Figure 4). No single OTU spans more than one plant division on Figure 4 (which describes plant genomic EVE contents). Fernendovirus 1 sequences were found in both fern genomes and a lycopod transcriptome, but were found in no other genomes. Gymnosperm genomes include exclusively ECRT loci that are assigned to one of the four Gymnendovirus OTUs, all of which were undetected outside of gymnosperms. Among gymnosperms, the three conifer genomes analyzed contain a mixture ECRTs from the four Gymnendovirus OTUs. By contrast, only ECRT loci classified as Gymnendovirus 2 were detected in *Ginkgo biloba* (*Ginkgoales*). Within angiosperms, we also observed a dichotomy for the distribution of ECRTs between monocots and dicots. On one hand, Yendovirus, *Badnavirus*, Orendovirus and Florendovirus ECRTs are common in monocots, with Orendovirus ECRTs being the only monocot-specific ones. On the other hand, *Petuvirus*, Florendovirus, Xendovirus, *Cavemovirus*/*Solendovirus* and Yendovirus ECRTs are most widely distributed in dicots, Florendovirus and Yendovirus hence being remarkably well represented in both dicots and monocots.

## Discussion

Endogenous viral elements are considered relics of past infections, and an extrapolation of the results from this study is that nearly every tracheophyte plant species in the world has at some point in its evolutionary history been subject to infection by at least one, and sometimes five distinct viral species/genera from the family *Caulimoviridae*. This finding attests to the tremendous adaptability of the *Caulimoviridae*, surpassing any other group of plant viruses. Members of the *Caulimoviridae* have likely also had a large influence on plant evolution, either as pathogens or as donors of novel genetic material to the plant genome.

A defining moment in the evolution of the *Caulimoviridae* appears to be the development of vasculature in plants. The presence of a 30K movement protein is an important feature of the *Caulimoviridae* that distinguishes it from the LTR retrotransposon family *Metaviridae*, and this protein is crucial for the formation of systemic infection by allowing intercellular trafficking of macromolecules through increasing the size exclusion limit of plasmodesmata ^21^. Although algae contain plasmodesmata, which superficially resemble those of higher plants, they are not homologous structures ^22^. While the acquisition of a 30K movement protein would have provided a selective advantage for ancestral caulimovirids to colonize the tracheophytes, it would not have facilitated infection of more primitive plant forms.

When recapitulating the distribution of EVEs in plant genomes, the known host range of exogenous viruses, and the phylogenetic relationships between caulimovirid OTUs and major groups of vascular plants (Figure 5) to infer the evolutionary trajectories of plant-virus coevolution, we obtain a complex pattern of host-virus associations. At the OTU level, the host distributions of petu- and xendovirus, including dicots and the ANITA grade angiosperm (*Amborella trichopoda*) but not any of the monocot species, is suggestive of horizontal transfer. In addition, although vertical transmission is overall well supported by a co-evolutionary study of florendoviral EVEs and their host species ^10^, it could not be confirmed for *A. trichopoda*. Together with the observation that *A. trichopoda* presents a high density of ECRTs (Figure 2B), this may suggest that this species is permissive to infection by a range of caulimovirid genera and/or that it represents a hotbed for the emergence of caulimovirid genera, some of which swapped towards mesangiosperms.

**Figure 5.**
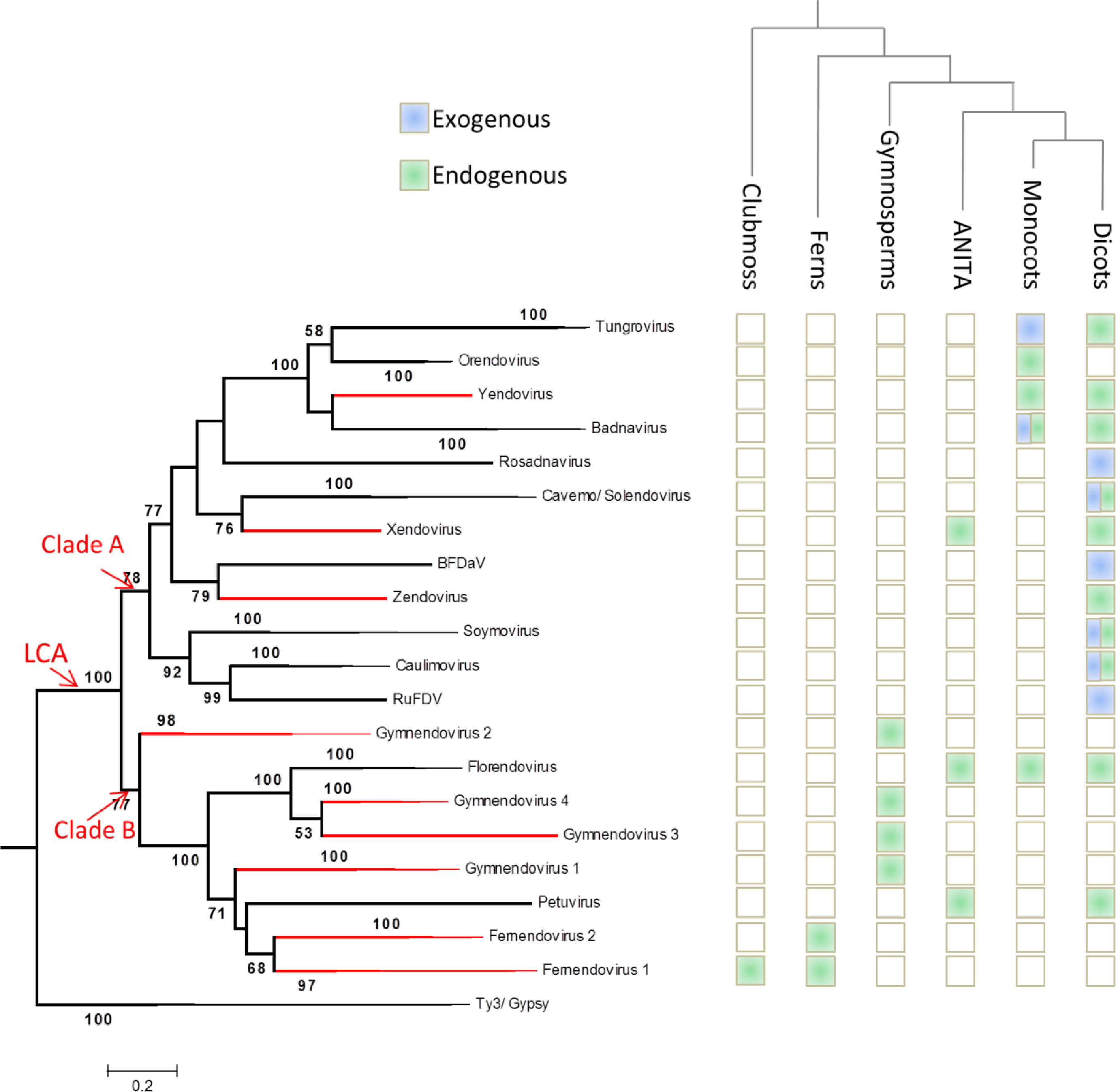
Working scenario of *Caulimoviridae* deep evolution. The left tree is the same as in Figure 3 where the deepest *Caulimoviridae* node was annotated as LCA (last common ancestor). The top cladogram indicates the evolutionary relationships between major classes of vascular plants. At the intersection between both trees, color code indicates the presence of EVEs (green) and of known associations with exogenous viruses (blue).

At a deeper level within the caulimovirid tree, clade A caulimovirids were found exclusively in mesangiosperm species. Clade B viruses, instead, were found to associate with plants from all the major classes of Tracheophyta. Assuming the monophyly of clades A and B, the obtained plant-virus associations could be explained by the emergence of these viruses in a common ancestor of the gymnosperms and angiosperms followed by several major host swaps: in clade A, between monocots and dicots, and in clade B between gymnosperms and angiosperms in the case of florendoviruses and petuviruses (although the position of this latter genus in the tree is uncertain), and from gymnosperms to ferns and clubmoss in the case of fernendoviruses. Following this scenario, the *Caulimoviridae* could have emerged at the latest with the Spermatophyta, *i.e*. during the Devonian era, about 320 MYA ^23^. Such a scenario would however imply several host swaps that overlay plant evolutionary history and incomplete sampling in clade A as the OTUs that associate with monocots are not sister to but nested in those that associate with dicots. We therefore do not rule out the possibility that the observed plant-virus associations reflect to some extent stochastic survival (or sampling) of a diversity of ancestral members of the *Caulimoviridae* that would have arose at latest during the emergence of clubmoss towards the end of the Silurian era 420 MYA.

## Methods

### Discovery and clustering of novel Caulimoviridae OTUs

We built a library containing an assortment of amino acid (aa) sequences from 54 RT domains including four from *Retroviridae*, six from Ty3/Gypsy LTR retrotransposons, 41 from eight different *Caulimoviridae* genera (Florendovirus, *Caulimovirus*, *Tungrovirus*, *Cavemovirus*, *Solendovirus*, *Badnavirus*, *Soymovirus*, and *Petuvirus*), two from *Picea glauca*, and the one from the DIRS-1 element. We compared this library to a collection of 72 genome assemblies from the *Viridiplantae* (listed in Supplementary Table 1) using tBLASTn with default parameters (except –e=1e-5). The hit genomic loci were merged when overlapping and their coordinates were extended 120 bases upstream and downstream. Extended hit loci were translated and the protein sequences of length >=200aa were compared to the initial RT library using BLASTp with default parameters (except – e=1e-5). Queries with best alignment score against Caulimoviridae over at least 170 residues were selected for further analysis. For each plant species, the selected set of RT aa sequences have been clustered following sequence similarity using the UCLUST program ^24^ with identity threshold set at 80%. The longest sequence from each resulting cluster was considered as the representative sequence and it was appended to the initial RT library. To detect potential false positives, each set of sequences (each consisting of the initial RT library and cluster representatives from one species) was aligned using MUSCLE followed by filtering of lower fit sequences using two rounds of trimAl v1.2 ^25^ to remove poorly aligned sequences (-resoverlap 0.75 -seqoverlap 50) separated by one round to remove gaps from the alignment (-gt 0.5). The representative sequences from each plant species that passed this selection were combined into a single file and appended to the initial RT library to be clustered with UCLUST using an identity threshold of 55%. At this level of similarity, aa RT sequences from every genus in the *Caulimoviridae* is separated except those from *Cavemovirus* and *Solendovirus*.

Starting with the first cluster, one or more sequences presenting high quality alignment and containing several conserved residues as determined contextually for each cluster were then manually selected to be representative of the diversity observed within each cluster. The following clusters were processed similarly while keeping the representative sequences selected from previously processed clusters. Clusters containing ECRT sequences from only one plant species were analyzed only when they contained at least three sequences. After processing each cluster individually, a total of 56 ECRT sequences detected here and 20 RT from known genera have been selected for their remarkable divergence. Together with four RT sequences from Ty3/Gypsy LTR retrotransposons, these combined sequences (hereafter referred to as “diverse library”) were aligned with the GUIDANCE2 ^26^ program using MAFFT ^27^ to generate bootstrap supported MSA and to remove columns (–colCutoff) with confidence score below 0.95 (16/244 columns removed in the RT sequence from Caulimovirus CaMV). The resulting MSA was then used to build the phylogenetic network shown in Figure 1 and Supplementary Figure 1 with SplitsTree4 ^28^ applying the NeighborNet method with uncorrected P distance model and 1,000 bootstrap tests. Manual analysis of this network enabled the discrimination of 17 distinct groups sharing deep connections among caulimovirid sequences.

In response to the discovery of several novel OTUs, we repeated ECRT mining in plant genomes using the diverse library as query. This second search is also designed to be more sensitive as it takes into account DNA sequences instead of uninterrupted ORFs. The workflow is identical to the one employed for the initial search until obtaining the set of extended hit loci. These were directly compared to the diverse library using BLASTx with default parameters (except –e=1e-5). Queries with best alignment score against any Caulimoviridae with an alignment length above 80% of subject length (set generically to 576 bp considering an average size of RT domains of 240 aa) were selected for phylogenetic placement.

### Phylogenetic analysis

Fragments of virus sequence were assembled using CodonCode aligner 6.0.2 using default settings or using VECTOR NTI Advance 10.3.1 (Invitrogen) operated using default settings, except that the values for maximum clearance for error rate and maximum gap length were increased to 500 and 200, respectively.

Phylogenetic reconstruction was performed using the contiguous nucleotide sequences corresponding to the protease, reverse transcriptase and ribonuclease H domains. Whole sequences from Caulimoviridae genera representatives and Ty3 and Gypsy LTR retrotransposons were first aligned with global method using MAFFT v7.3 ^27^. The core genomes was extracted and re-aligned by local method using MAFFT. The resulting alignment was tested for different evolutionary models with pmodeltest v1.4 (from ETE 3 package ^29^) which inferred the GTRGAMMA model. Phylogenetic inference with maximum likelihood was then performed using RaxML v8.2 ^30^ under the predicted model with 500 ML bootstrap replicates.

The resulting tree was then used as a reference to classify the ECRT loci mined from plant genomes. We first added query sequences from each plant species separately to the reference alignment and aligned each library using Mafft v7.3 (with options –addfragment, –keeplength and by reordering). We then tested the most likely placement of each ECRT sequence on to the reference tree using pplacer v1.1 alpha19 ^20^ with the option (–keep-at-most 1) with allows to keep one placement for each query sequence. The python package Taxit was used to construct a reference package which we used to run pplacer.

### Data availability

The datasets and scripts generated during during the current study are available from the corresponding author on request.

## Acknowledgements

We thank Mark Tepfer for critical reading of a previous version of this manuscript. PYT was funded by the Guadeloupe Region and the European Regional Development Fund.

## Author contributions

SD, ADW, and FM performed research; FAD and ML provided research environment; SD, ADW, PYT, and FM analyzed the data; ADW, PYT, and FM wrote the manuscript with contributions from SD; FM designed the research.

## Supplementary material

Supplementary file 1: reconstructed ancestral sequences of members of the *Caulimoviridae* from novel OTUs (fasta format) and onekp contigs.

**Supplementary Figure 1:**
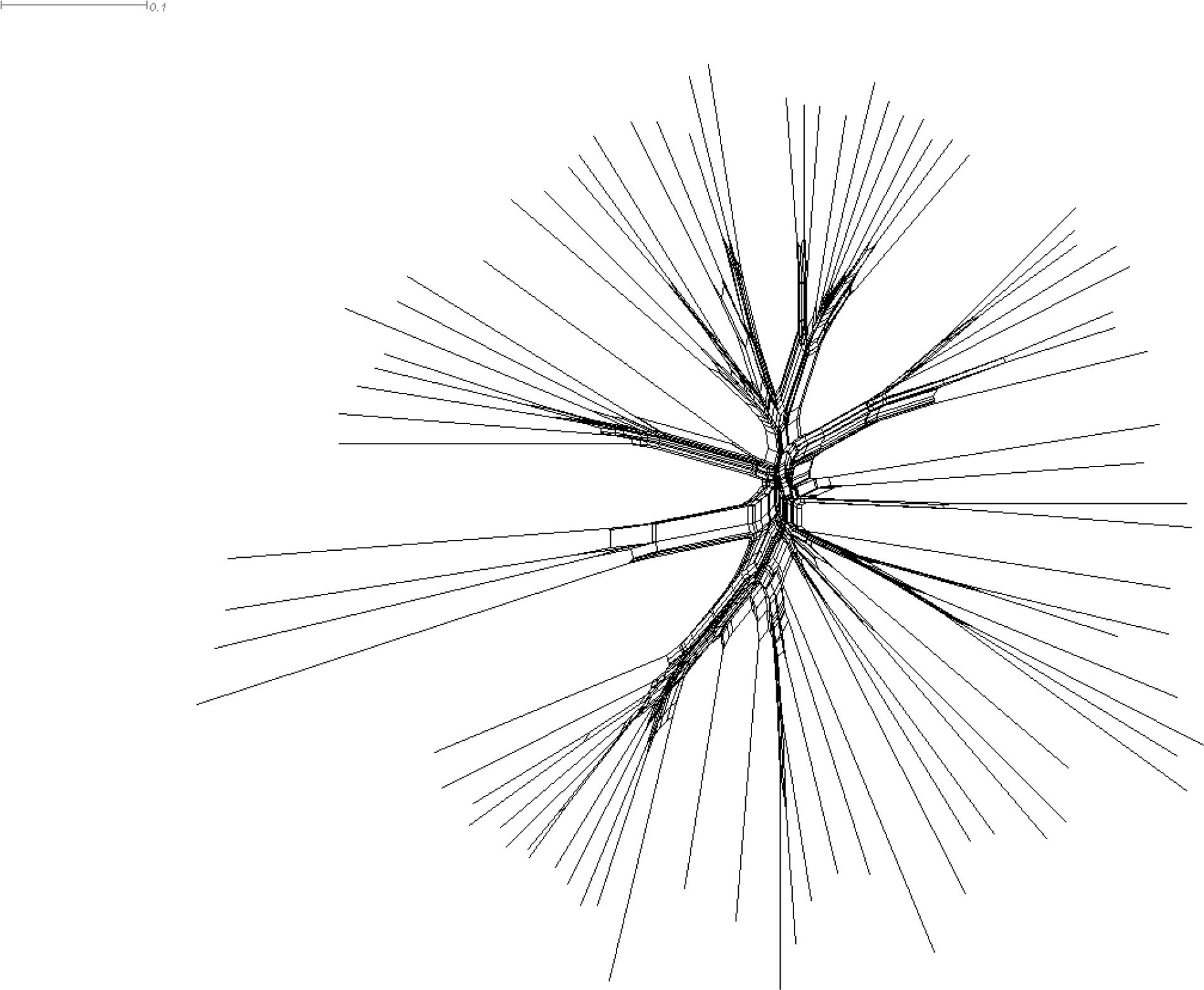
Overview of the phylogenetic network used to build Figure 1.

**Supplementary Figure 2:**
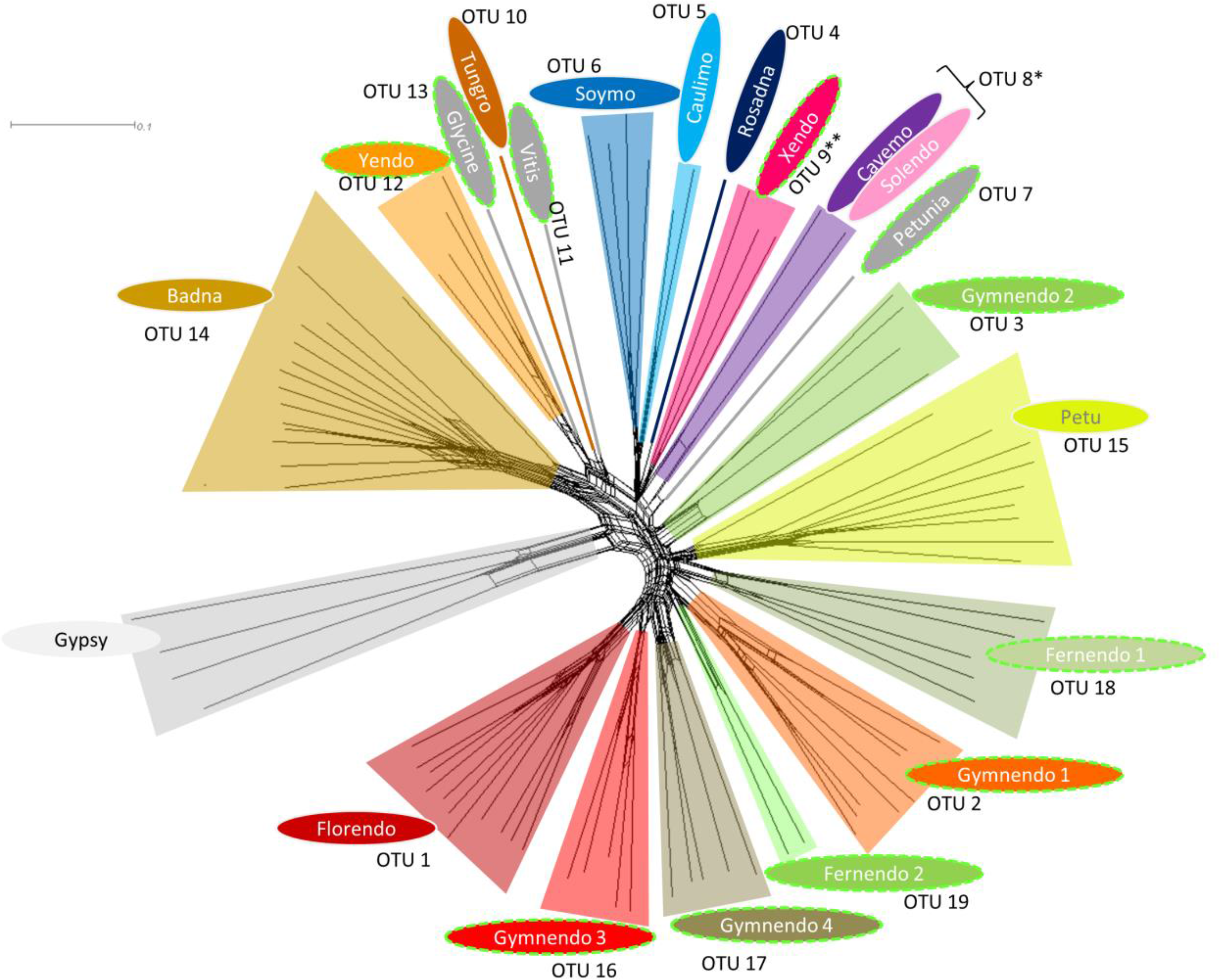
ECRT ORFs collected from ferns cluster as two novel OTUs. Representative sequences identified in fern genomes were appended to the collection of sequences represented in Figure 1. The resulting library has been re-aligned with MUSCLE and phylogenetic network was built using SplitsTree. The branches containing fern sequences have been empirically grouped into two novel OTUs (OTU 18 and OTU 19).

